## Supplementary material for "Can education be personalised using pupils’ genetic data?"

We split educational achievement and the polygenic scores by quintiles and deciles to assess the level of agreement (Table S1). The agreement between the quantiled measures of achievement at ages 7 and 16 and the polygenic scores were slightly higher than expected. For the GWAS significant polygenic score, the Kappa statistics show that agreement was at most only 5% higher than would be expected by random agreement compared to perfect agreement (quintiles $\kappa$ = 0.05 for achievement at age 7). Agreement was higher for the all SNP polygenic score and generally higher for age 16 than age 7 achievement. At age 16, agreement was at least twice as high for quantiles of prior achievement than the polygenic scores when compared to random allocation. Agreement with age 16 achievement was highest for age 14 achievement, being 46% and 28% better than expected by chance for quintiles and deciles of achievement respectively.

Table S1: Agreement between educational achievement (EA) quantiles and other quantiled measures. Polygenic scores (PGS) built using only genome-wide significant SNPs (GWAS sig SNPs) or all education associated SNPs (All SNP PGS) from the largest GWAS of educational attainment[1]. p values for all tests <0.001.

| | Age 7 achievement | | Age 16 achievement | |
|---|---|---|---|---|
| | Agreement | Kappa | Agreement | Kappa |
| **Quintiles** | | | | |
| *Genotypic predictor* | | | | |
| GWAS sig PGS | 24.1% | 0.05 (0.01) | 23.2% | 0.04 (0.01) |
| All SNPs PGS | 26.3% | 0.08 (0.01) | 28.2% | 0.10 (0.01) |
| *Phenotypic predictor* | | | | |
| Age 14 EA | | | 56.5% | 0.46 (0.01) |
| Age 11 EA | | | 46.8% | 0.33 (0.01) |
| Age 7 EA | | | 37.8% | 0.22 (0.01) |
| **Deciles** | | | | |
| *Genotypic predictor* | | | | |
| GWAS sig PGS | 11.4% | 0.02 (0.01) | 12.4% | 0.03 (0.01) |
| All SNPs PGS | 13.9% | 0.04 (0.01) | 14.3% | 0.05 (0.01) |
| *Phenotypic predictor* | | | | |
| Age 14 EA | | | 35.1% | 0.28 (0.01) |
| Age 11 EA | | | 27.5% | 0.19 (0.01) |
| Age 7 EA | | | 20.9% | 0.12 (0.01) |

**Table S2: Variance explained in educational** achievement **at age 7.** Polygenic scores (PGS) built using only genome-wide significant SNPs (GWAS sig SNPs) or all education associated SNPs (All SNP PGS) from the largest GWAS of educational attainment[1]. FSM, Free School Meals. EFL, English as a Foreign Language. SEN, Special Educational Needs. Eduyears, years of education. SEP, socioeconomic position.

| Polygenic score (PGS) | Covariates | R2 | Lower 95% CI from bootstraps | Upper 95% CI from bootstraps |
|---|---|---|---|---|
| No PGS | Age, sex | 0.060482 | 0.043601 | 0.077363 |
| GWAS sig PGS | Age, sex | 0.083937 | 0.066095 | 0.101779 |
| All SNP PGS | Age, sex | 0.123893 | 0.102153 | 0.145633 |
| No PGS | Age, sex, FSM, EFL, SEN | 0.167781 | 0.142809 | 0.192753 |
| GWAS sig PGS | Age, sex, FSM, EFL, SEN | 0.185696 | 0.160279 | 0.211113 |
| All SNP PGS | Age, sex, FSM, EFL, SEN | 0.215013 | 0.189151 | 0.240875 |
| No PGS | Age, sex, FSM, EFL, SEN, parents Eduyears | 0.238826 | 0.213266 | 0.264386 |
| GWAS sig PGS | Age, sex, FSM, EFL, SEN, parents Eduyears | 0.247198 | 0.221092 | 0.273303 |
| All SNP PGS | Age, sex, FSM, EFL, SEN, parents Eduyears | 0.257594 | 0.230686 | 0.284502 |
| No PGS | Age, sex, FSM, EFL, SEN, parents Eduyears, parents SEP | 0.244971 | 0.216829 | 0.273113 |
| GWAS sig PGS | Age, sex, FSM, EFL, SEN, parents Eduyears, parents SEP | 0.252427 | 0.22521 | 0.279644 |
| All SNP PGS | Age, sex, FSM, EFL, SEN, parents Eduyears, parents SEP | 0.262971 | 0.234203 | 0.291738 |

**Table S3: Incremental R2 for educational achievement at age 7.** Polygenic scores (PGS) built using only genome-wide significant SNPs (GWAS sig SNPs) or all education associated SNPs (All SNP PGS) from the largest GWAS of educational attainment[1]. FSM, Free School Meals. EFL, English as a Foreign Language. SEN, Special Educational Needs. Eduyears, years of education. SEP, socioeconomic position.

| Polygenic score (PGS) | Covariates | Incremental gain in R2 | Lower 95% CI from bootstraps | Upper 95% CI from bootstraps |
|---|---|---|---|---|
| GWAS sig PGS | Age, sex | 0.023455 | 0.005614 | 0.041297 |
| All SNP PGS | Age, sex | 0.063411 | 0.041671 | 0.085151 |
| GWAS sig PGS | Age, sex, FSM, EFL, SEN | 0.017915 | -0.0075 | 0.043332 |
| All SNP PGS | Age, sex, FSM, EFL, SEN | 0.047232 | 0.02137 | 0.073094 |
| GWAS sig PGS | Age, sex, FSM, EFL, SEN, parents Eduyears | 0.008372 | -0.01773 | 0.034477 |
| All SNP PGS | Age, sex, FSM, EFL, SEN, parents Eduyears | 0.018768 | -0.00814 | 0.045676 |
| GWAS sig PGS | Age, sex, FSM, EFL, SEN, parents Eduyears, parents SEP | 0.007456 | -0.01976 | 0.034673 |
| All SNP PGS | Age, sex, FSM, EFL, SEN, parents Eduyears, parents SEP | 0.018 | -0.01077 | 0.046767 |

**Table S4: Variance explained in educational** achievement **at age 16.** Polygenic scores (PGS) built using only genome-wide significant SNPs (GWAS sig SNPs) or all education associated SNPs (All SNP PGS) from the largest GWAS of educational attainment[1]. FSM, Free School Meals. EFL, English as a Foreign Language. SEN, Special Educational Needs. EA7, prior educational achievement at age 7. EA11, prior educational achievement at age 11. EA14, prior educational achievement at age 14. Eduyears, years of education. SEP, socioeconomic position.

| Polygenic score (PGS) | Covariates | R2 | Lower 95% CI from bootstraps | Upper 95% CI from bootstraps |
|---|---|---|---|---|
| No PGS | Age, sex, | 0.032907 | 0.019845 | 0.04597 |
| GWAS sig PGS | Age, sex, | 0.066455 | 0.049561 | 0.083349 |
| All SNP PGS | Age, sex, | 0.16226 | 0.138444 | 0.186076 |
| No PGS | Age, sex, FSM, EFL, SEN | 0.159373 | 0.133462 | 0.185284 |
| GWAS sig PGS | Age, sex, FSM, EFL, SEN | 0.18515 | 0.15853 | 0.21177 |
| All SNP PGS | Age, sex, FSM, EFL, SEN | 0.262899 | 0.235065 | 0.290733 |
| No PGS | Age, sex, FSM, EFL, SEN, EA7 | 0.465058 | 0.439548 | 0.490569 |
| GWAS sig PGS | Age, sex, FSM, EFL, SEN, EA7 | 0.471385 | 0.446809 | 0.495962 |
| All SNP PGS | Age, sex, FSM, EFL, SEN, EA7 | 0.503215 | 0.479072 | 0.527359 |
| No PGS | Age, sex, FSM, EFL, SEN, EA11 | 0.501463 | 0.454133 | 0.548793 |
| GWAS sig PGS | Age, sex, FSM, EFL, SEN, EA11 | 0.508429 | 0.464163 | 0.552694 |
| All SNP PGS | Age, sex, FSM, EFL, SEN, EA11 | 0.534566 | 0.493807 | 0.575325 |
| No PGS | Age, sex, FSM, EFL, SEN, EA14 | 0.651496 | 0.608533 | 0.694458 |
| GWAS sig PGS | Age, sex, FSM, EFL, SEN, EA14 | 0.653617 | 0.611231 | 0.696003 |
| All SNP PGS | Age, sex, FSM, EFL, SEN, EA14 | 0.663347 | 0.622094 | 0.704601 |
| No PGS | Age, sex, FSM, EFL, SEN, EA7, EA11, EA14 | 0.676328 | 0.64597 | 0.706686 |
| GWAS sig PGS | Age, sex, FSM, EFL, SEN, EA7, EA11, EA14 | 0.677808 | 0.647664 | 0.707951 |
| All SNP PGS | Age, sex, FSM, EFL, SEN, EA7, EA11, EA14 | 0.686275 | 0.656997 | 0.715552 |

**Table S5: Incremental R2 for educational achievement at age 16**. Polygenic scores (PGS) built using only genome-wide significant SNPs (GWAS sig SNPs) or all education associated SNPs (All SNP PGS) from the largest GWAS of educational attainment[1]. FSM, Free School Meals. EFL, English as a Foreign Language. SEN, Special Educational Needs. EA7, prior educational achievement at age 7. EA11, prior educational achievement at age 11. EA14, prior educational achievement at age 14.

| Polygenic score (PGS) | Covariates | Incremental gain in R2 | Lower 95% CI from bootstraps | Upper 95% CI from bootstraps |
|---|---|---|---|---|
| Parents Eduyears | Age, sex, | 0.190153 | 0.165965 | 0.214341 |
| Parents SEP | Age, sex, | 0.213652 | 0.18835 | 0.238954 |
| GWAS sig PGS | Age, sex, | 0.033548 | 0.016653 | 0.050442 |
| All SNP PGS | Age, sex, | 0.129353 | 0.105536 | 0.153169 |
| GWAS sig PGS | Age, sex, FSM, EFL, SEN | 0.025778 | -0.00084 | 0.052398 |
| All SNP PGS | Age, sex, FSM, EFL, SEN | 0.103526 | 0.075692 | 0.13136 |
| GWAS sig PGS | Age, sex, FSM, EFL, SEN, EA7 | 0.006327 | -0.01825 | 0.030903 |
| All SNP PGS | Age, sex, FSM, EFL, SEN, EA7 | 0.038157 | 0.014014 | 0.062301 |
| GWAS sig PGS | Age, sex, FSM, EFL, SEN, EA11 | 0.006966 | -0.0373 | 0.051232 |
| All SNP PGS | Age, sex, FSM, EFL, SEN, EA11 | 0.033104 | -0.00766 | 0.073862 |
| GWAS sig PGS | Age, sex, FSM, EFL, SEN, EA14 | 0.002122 | -0.04026 | 0.044508 |
| All SNP PGS | Age, sex, FSM, EFL, SEN, EA14 | 0.011852 | -0.0294 | 0.053105 |
| GWAS sig PGS | Age, sex, FSM, EFL, SEN, EA7, EA11, EA14 | 0.00148 | -0.02866 | 0.031623 |
| All SNP PGS | Age, sex, FSM, EFL, SEN, EA7, EA11, EA14 | 0.009947 | -0.01933 | 0.039224 |

**Figure S1: Distributions of prior achievement between "high achievers" at age 16 and all other pupils.** Panel A: Prior achievement at age 7. Panel B: Prior achievement at age 11. Panel C: Prior achievement at age 14. Educational achievement (EA) measured using fine graded point scores from educational exams taken at ages 7, 11 and 14. s High achievers defined as pupils with age 16 educational exam scores in the top 10% of the sample. Prior attainment measured as educational achievement (EA) at ages.

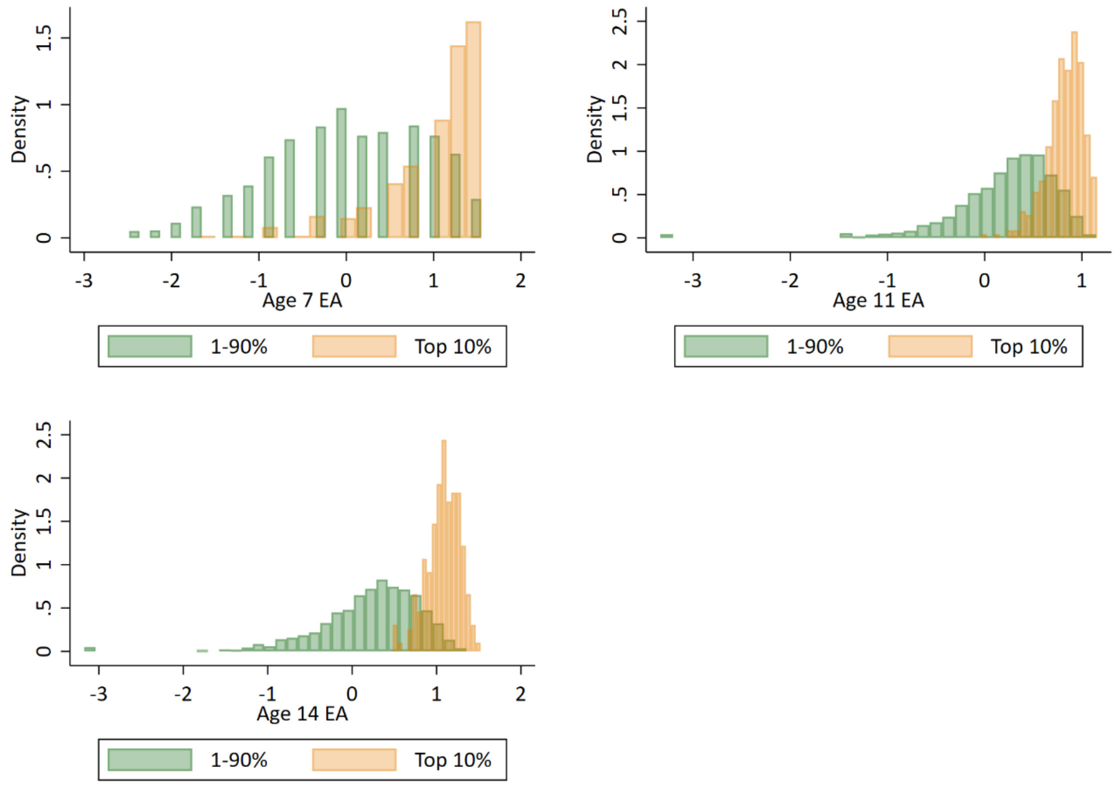

**Figure S2: Independent scatter plots showing correlation between realised achievement at age 16 and achievement predicted from variables with top 10% of predicted achievers highlighted in green.**
Educational achievement measured using fine graded point scores from educational exams. Predicted achievement at age 16 generated from a polygenic score built using all education associated SNPs (All SNP PGS) from the largest GWAS of educational attainment[1]. Parental educational attainment measured as average years of completed education. Parental socioeconomic position (SEP) measured as highest parental score on the Cambridge Social Stratification Score scale. All PGS analyses include adjustment for the first 20 principal components of population stratification.

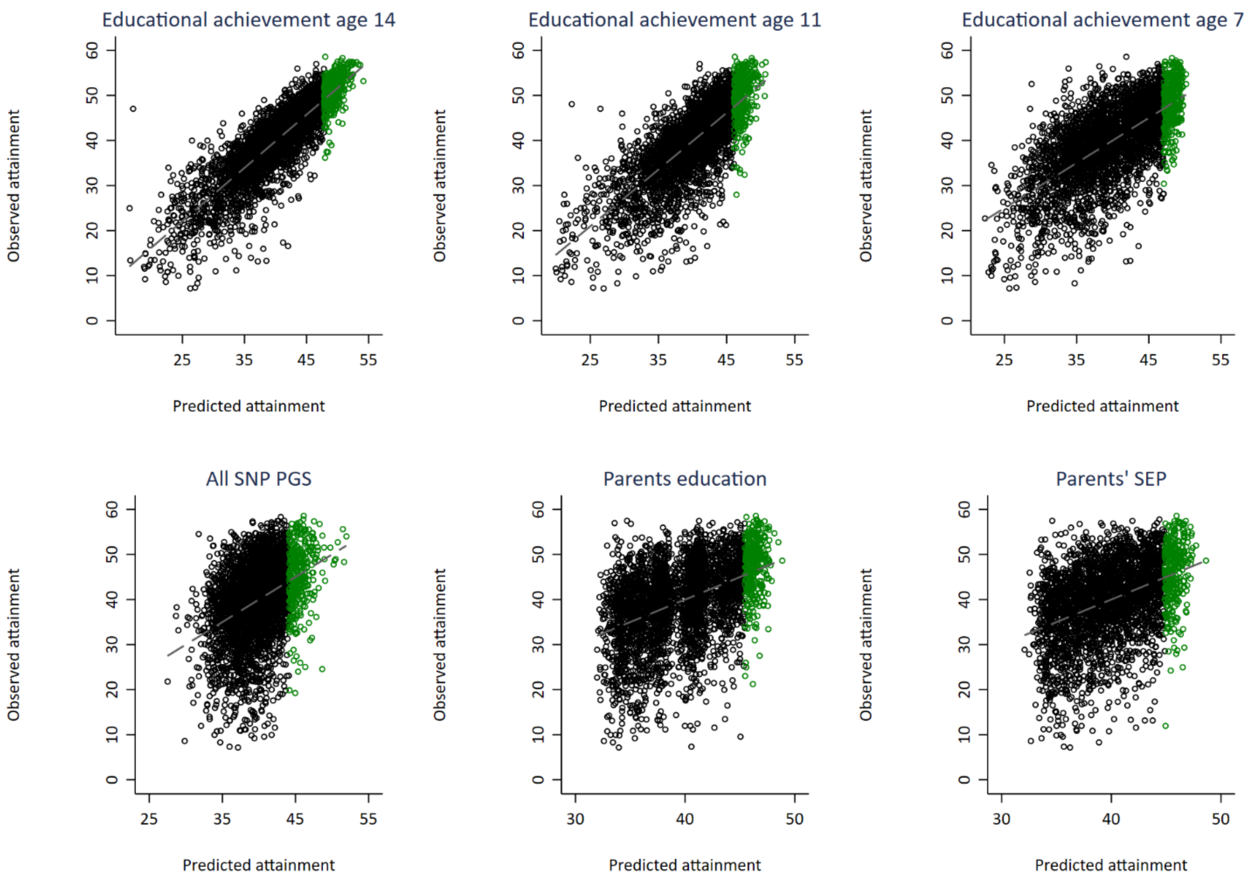

**Figure S3: Independent ROC curves for deciled measures of polygenic scores, parental education and parental socioeconomic position predicting high achieving students at age 7 defined at different thresholds (e.g. top 10%).** Parents educational attainment (EA) measured as years of completed education. Parents socioeconomic position (SEP) measured as highest parental score on the CAMSIS social class scale. Polygenic scores (PGS) built using only genome-wide significant SNPs (GWAS sig SNPs) or all education associated SNPs (All SNP PGS) from the largest GWAS of educational attainment[1]. All PGS analyses include adjustment for the first 20 principal components of population stratification. Note that x axis displays 1-specificty. Note that cut-offs of 1% and 5% could not be used due to lack of data richness in the age 7 achievement scores.

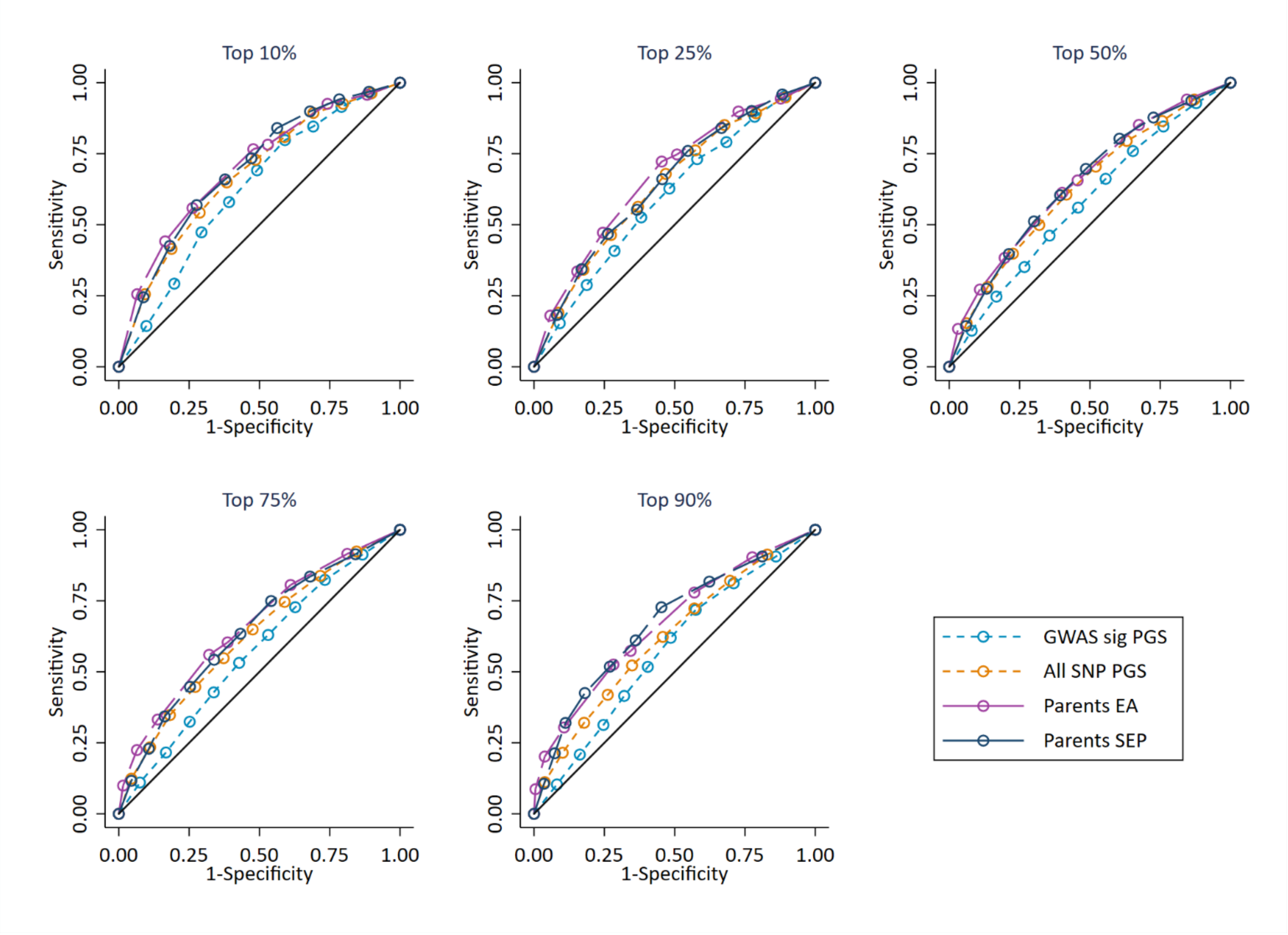

**Figure S4: ROC curves for deciled measures of polygenic scores predicting high achieving students at age 16 (pupils with age 16 educational exam scores in the top 10% of the sample) conditional on prior attainment and pupil characteristics.** Polygenic scores (PGS) built using only genome-wide significant SNPs (GWAS sig SNPs) or all education associated SNPs (All SNP PGS) from the largest GWAS of educational attainment[1]. Pupil characteristics include Free School Meals (FSM), English as a Foreign language (EFL) and Special Educational Needs (SEN) status. All analyses include adjustment for the first 20 principal components of population stratification. Note that x axis displays 1-specificty.

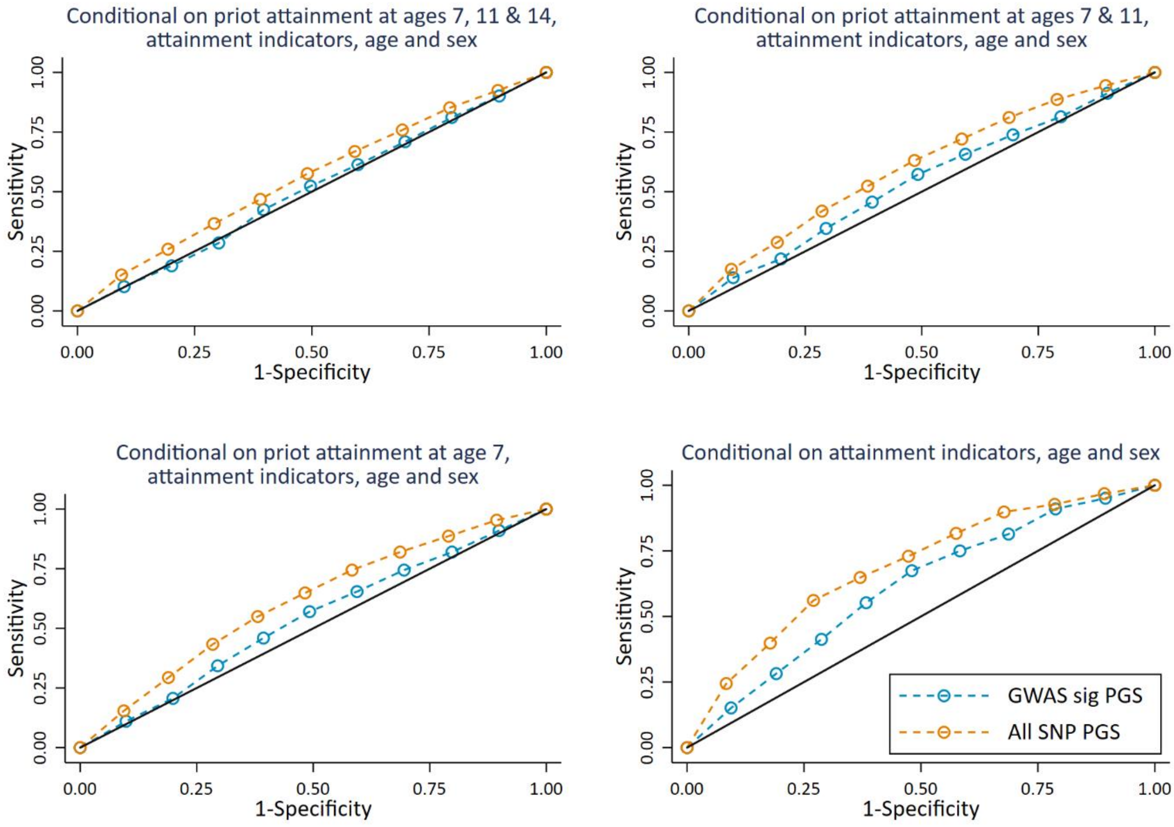

**Figure S5: Independent ROC curves for deciled measures of prior attainment and polygenic scores predicting high achieving students at age 16 defined at different thresholds (e.g. top 1%).** Polygenic scores (PGS) built using only genome-wide significant SNPs (GWAS sig SNPs) or all education associated SNPs (All SNP PGS) from the largest GWAS of educational attainment[1]. All PGS analyses include adjustment for the first 20 principal components of population stratification. Note that x axis displays 1-specificty.

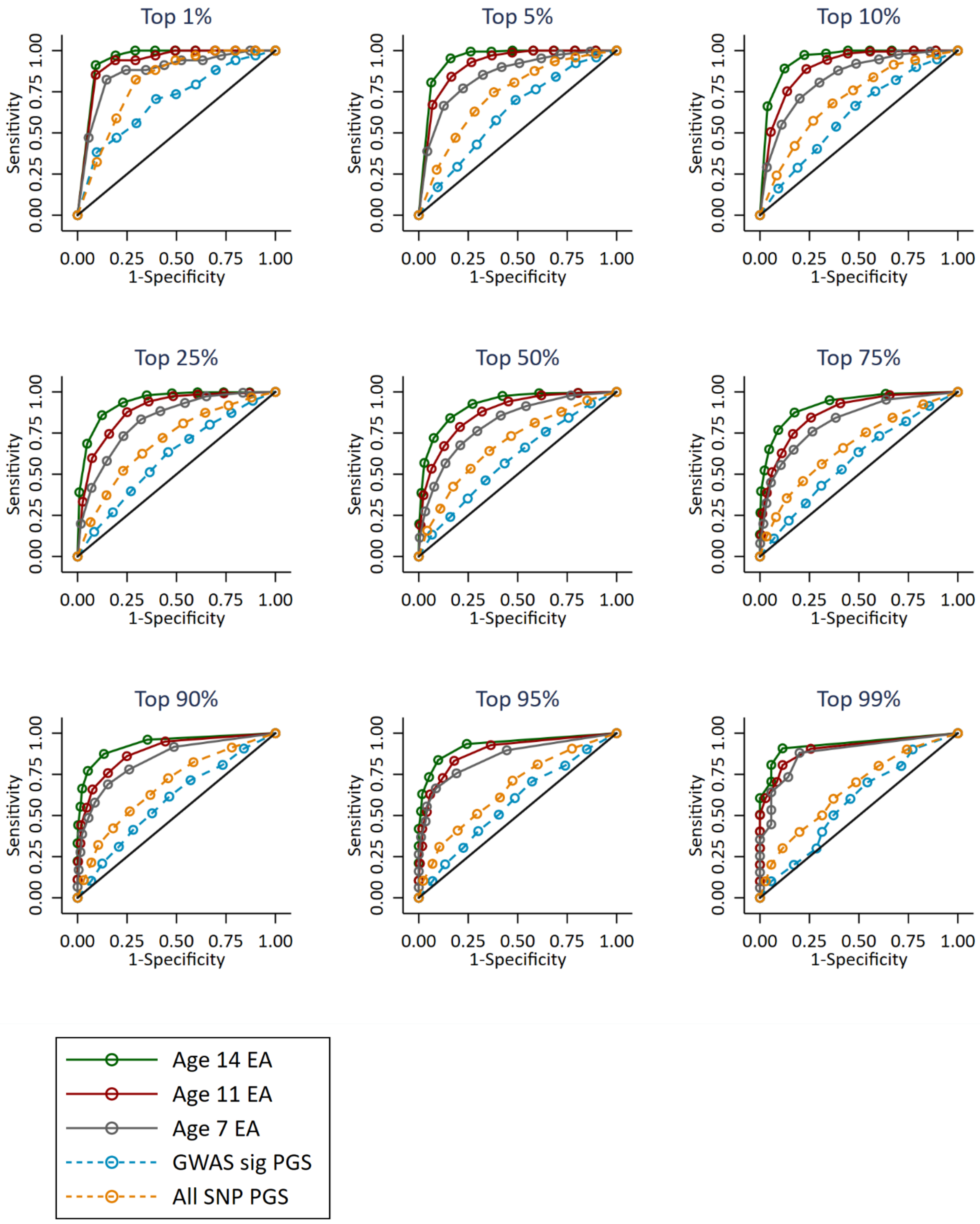

Figure S6: Attrition STROBE diagram for analytical sample.

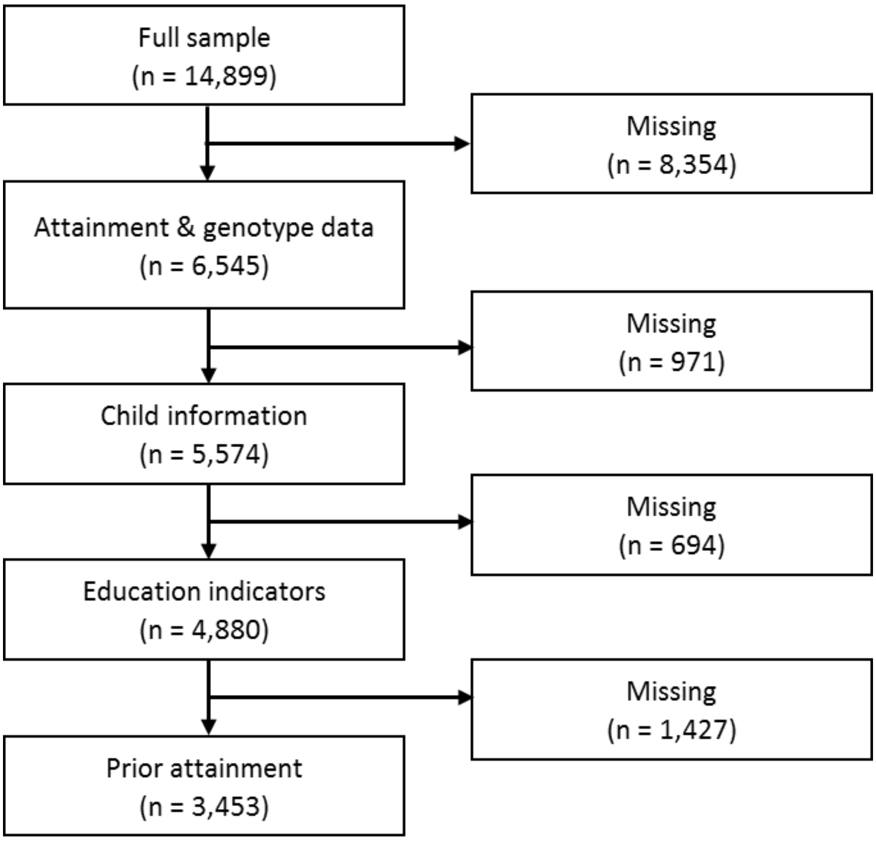
